# Real-time monitoring of cotranscriptional riboswitch folding and switching

**DOI:** 10.1101/2019.12.20.885053

**Authors:** Boyang Hua, Christopher P. Jones, Jaba Mitra, Peter J. Murray, Rebecca Rosenthal, Adrian R. Ferré-D’Amaré, Taekjip Ha

**Author notes:** Co-first authors (Boyang Hua, Christopher P. Jones).

## Abstract

Riboswitches function through cotranscriptional conformation switching governed by cognate ligand concentration, RNA folding and transcription elongation kinetics. To investigate how these parameters influence riboswitch folding, we developed a novel vectorial folding assay (VF) in which the superhelicase Rep-X sequentially liberates the RNA strand from a heteroduplex in a 5’-to-3’ direction, mimicking the nascent chain emergence during transcription. The RNA polymerase (RNAP)-free VF recapitulates the kinetically controlled cotranscriptional folding of a ZTP riboswitch, whose activation is favored by slower transcription, strategic pausing, or a weakened transcriptional terminator. New methods to observe positions and local rates of individual helicases show an average Rep-X unwinding rate similar to bacterial RNAP elongation (~60 nt/s). Real-time single-molecule monitoring captured folding riboswitches in multiple states, including an intermediate responsible for delayed terminator formation. These methods allow observation of individual folding RNAs as they occupy distinct folding channels within the landscape that controls gene expression and showed that riboswitch fate control is encoded in its sequence and is readily interpreted by a directionally moving protein even in the absence of an RNA polymerase.

## Introduction

As gene expression regulatory elements, riboswitches are structurally divided into two domains—an “aptamer” domain sufficient for ligand binding and an “expression platform” domain regulating gene expression.(Garst et al., 2011; Jones and Ferré-D’Amaré, 2017; Serganov and Nudler, 2013) These two domains share an overlapping sequence making their formation mutually exclusive. Ligand binding stabilizes the aptamer-containing conformation and prevents the shared segment being used to form the alternative conformation containing the expression platform.(Garst et al., 2011; Jones and Ferré-D’Amaré, 2017; Serganov and Nudler, 2013) It has been proposed that cotranscriptional RNA folding creates a time window for aptamer domain folding and ligand binding, which spans from the complete synthesis of the aptamer domain (*i.e.*, the earliest time when ligands can bind) to the synthesis of enough of the expression platform to interfere with binding.(Wickiser et al., 2005a; Wickiser et al., 2005b; Zhang et al., 2010) In this regime, the gene-regulatory outcome would depend on the relative rates of transcription elongation, RNA folding, and ligand binding but systematic investigation of such kinetic control model is hampered by the lack of methods to vary these properties independently. As riboswitches have gained interest as drugs targets,(Howe et al., 2015) understanding their mechanisms is a natural step to inform these pursuits.

The *Fusobacterium ulcerans* ZTP riboswitch regulates gene expression by binding to 5-aminoimidazole-4-carboxamide riboside 5’-monophosphate and triphosphate (ZMP and ZTP, respectively), which are elevated during folate stress.(Bochner and Ames, 1982; Kim et al., 2015; Rohlman and Matthews, 1990) The ZTP riboswitch often regulates through a mechanism of kinetically controlled transcription termination. In the absence of ZMP, a transcription terminator hairpin or “terminator” (*i.e.*, the expression platform) forms and terminates transcription before the gene coding region (Fig. S1a). In the presence of ZMP, the aptamer-containing conformation is stabilized by ligand binding, preventing terminator formation and allowing transcription to complete (Fig. S1a).

This kinetic control model has been studied in several riboswitches using bulk and single-molecule methods to understand the relationship between ligand binding and RNA structure. These approaches are often preceded by the observation of a discrepancy between equilibrium ligand binding measurements and cotranscriptional riboswitch activation measurements. This discrepancy is then further examined through monitoring the ligand binding and RNA folding kinetics. For example, bulk approaches have made use of structural probing techniques, NMR spectroscopy and fluorescently labeled ligands to measure the RNA folding and ligand binding kinetics for different lengths of riboswitch RNAs representing different points in transcription.(Helmling et al., 2017; Wickiser et al., 2005a; Wickiser et al., 2005b) More recently, chemical probing approaches were combined with high-throughput sequencing and used to examine riboswitch structure during “roadblocked” transcription to obtain more complete maps of RNA folding for all transcript lengths.(Strobel et al., 2019; Watters et al., 2016) However, these approaches cannot measure real time folding dynamics because RNA conformations are measured long after transcription is stalled by a blockade.

Using an elegant optical tweezer assay, Frieda and Block observed the folding process of an adenine riboswitch during transcription.(Frieda and Block, 2012) Supporting the kinetic control model, the adenine aptamer was shown to fold and bind adenine no more than once before transcription decision. However, due to technical limitations, the roles of several important parameters, such as elongation speed and pausing, were not tested. In addition, both a drawback and an advantage of the optical tweezer assay are that tension is applied to either the nucleic acid substrates or RNAP, thus allowing for force-dependent perturbation of the system.

To date, experiments studying the kinetic control model often use RNAP (*i.e.*, T7 RNA polymerase or *Escherichia coli* RNAP holoenzyme). Assembly of RNAP on labeled nucleotide substrates enables the study of RNAP by single molecule FRET(Uhm et al., 2018) but restricts dye placement to the 5’ end of the RNA and requires partial transcription of the RNA. Another issue that arises when using RNAP to examine cotranscriptional RNA folding is that RNA mutations or experimental changes can affect both RNA folding and interactions between RNAP and nucleic acid substrates. Due to the complex behavior of RNAP, we sought an alternative for examining cotranscriptional RNA folding that would allow individual transcription components to be isolated and assayed. Below, we have used the Rep-X helicase(Arslan et al., 2015; Hua et al., 2018a; Mitra and Ha, 2019a) to mimic sequential (or vectorial) RNA folding without additional polymerase characteristics or limitations in dye placement. This assay also allowed us to tune the RNA release speed and to introduce pausing at arbitrary positions, which may also remodel the RNA folding landscape. We also report new assays to monitor the speed of the helicase and therefore the rate at which a nascent RNA is exposed to solution.

## Results

### Comparing aptamer domain versus full-length riboswitch folding

Here, we examined the *F. ulcerans* ZTP riboswitch (Fig. 1a, S1a) using single-molecule and bulk assays. The 75-nt riboswitch aptamer lacking the terminator sequence (Δterm) binds ZMP with an apparent dissociation constant (*K*_d_) of ~0.5 μM.(Jones and Ferré-D’Amaré, 2015) However, ZMP binding is sensitive to 3’-end extension after nt 75, and variants containing as few as 5 base pairs of the terminator bind poorly to ZMP (Fig. S1b). These measurements suggest that, at equilibrium, there should exist a window for ZMP to bind as the first ~10 nt of terminator sequence is being synthesized. We labeled Δterm with donor (Cy3) and acceptor (Cy5) fluorophores(Jones et al., 2019a) and tested for a ZMP-responsive conformational change using single-molecule FRET (Fig. 1a). When folded using a thermal refolding protocol (Online Methods), Δterm adopted a low FRET efficiency conformation (*E*_FRET_ ~0.25) in the absence of ZMP, and shifted to a mid-*E*_FRET_ conformation (~0.6) upon ZMP binding (Fig. 2a). In the absence of ZMP, Δterm predominantly occupied the low-*E*_FRET_ conformation but infrequently visited a mid-*E*_FRET_ conformation like the ZMP-bound state (*E*_FRET_ ~0.6, Fig. 2b), indicating transient conformational changes in the absence of ZMP. The *K*_d_ of the fluorophore-labeled Δterm to ZMP (1.3 μM, Fig. 2c) is only ~2.5-fold higher than that of the unlabeled Δterm (~0.5 μM),(Jones and Ferré-D’Amaré, 2015) suggesting that fluorophore labeling and surface attachment have only a modest effect on binding. At saturating ZMP (≥ 1 mM), ~30% of Δterm persisted in the low-*E*_FRET_ conformation (Fig. 2c), possibly due to a misfolded subpopulation incapable of ZMP binding.

**Figure 1.**
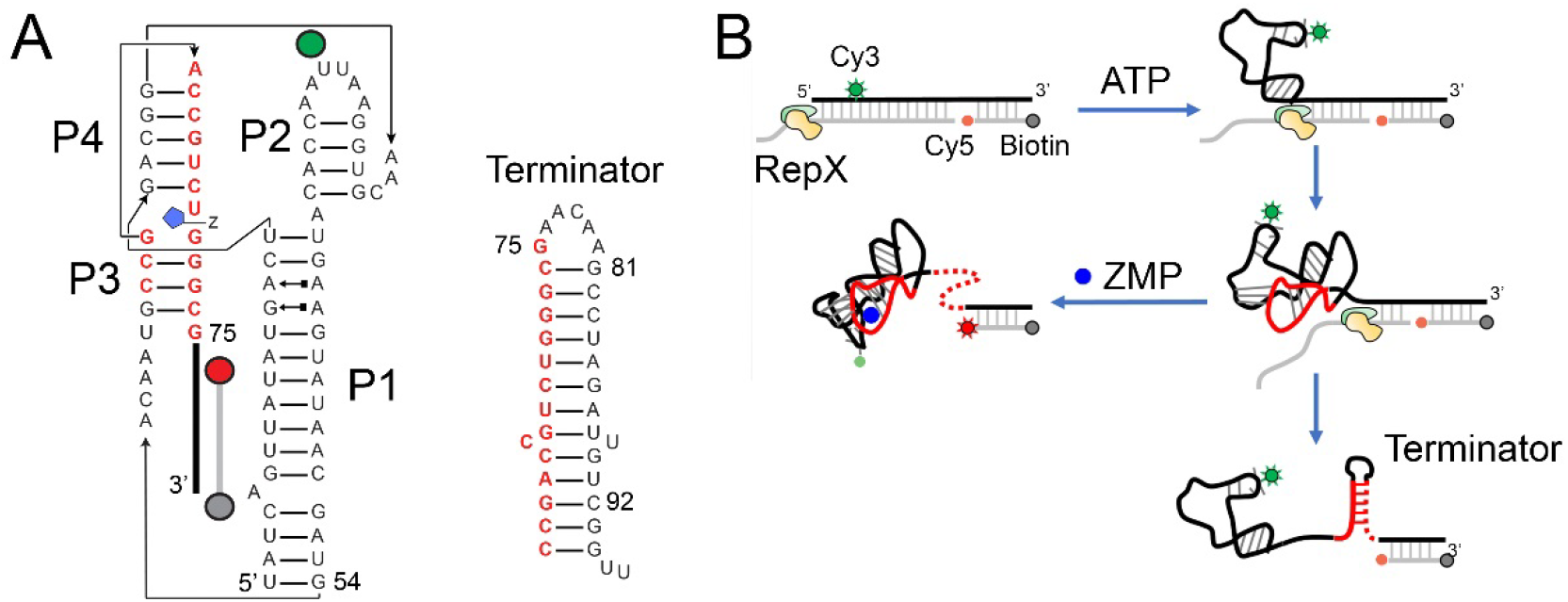
Single-molecule helicase-catalyzed unwinding experiments of the *F. ulcerans* ZTP riboswitch. (**a**) The ZTP riboswitch aptamer (Δterm, left) formed by paired elements P1-4. A Cy3 fluorophore (green sphere) is attached to solvent accessible U32, and a Cy5 fluorophore (red sphere) is placed at the 3’-end of the DNA tether, which is 5’-end labeled with biotin (grey sphere). In the full-length riboswitch (WT), the nucleotides colored in red switch between two conformations, one in which a terminator hairpin or “terminator” folds (right). (**b**) Cartoon depicting the VF assay, in which an engineered helicase Rep-X releases the riboswitch RNA (black) in a 5’-to-3’ direction by translocating on the cDNA strand (grey) and unwinding in a 3’-to-5’ direction.

**Figure 2.**
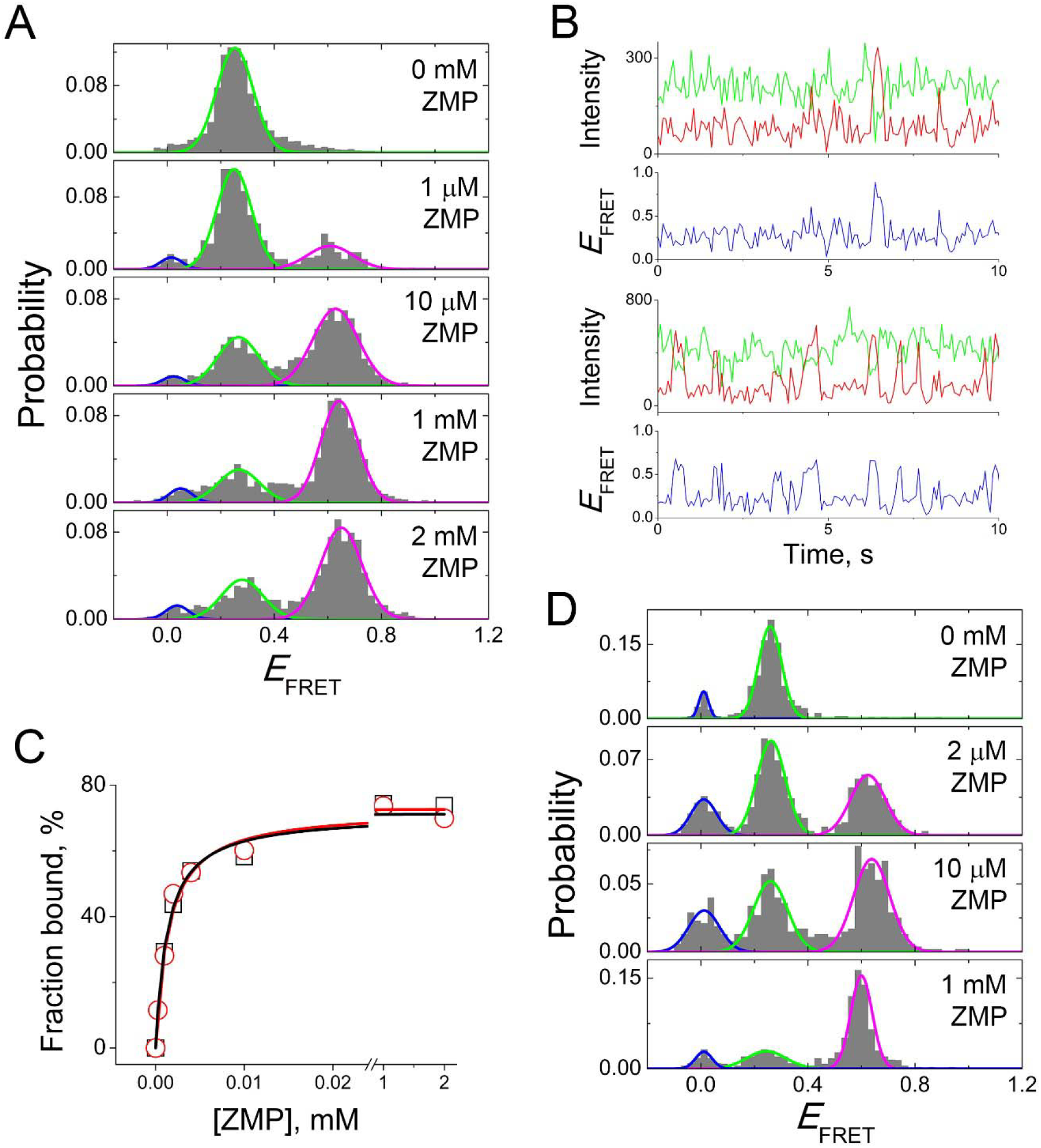
ZMP binding to the ZTP riboswitch aptamer after thermal refolding and vectorial folding. (**a**) The 75-nt ZTP riboswitch aptamer domain (Δterm) was thermally refolded at 0 mM ZMP and then incubated with different concentrations of ZMP. Gaussian fitting with global constraints was used to determine the relative population of the ZMP-bound (magenta) *vs.* ZMP-unbound (green) aptamer. (**b**) Example single molecule trajectories of thermally refolded Δterm in the absence of ZMP (top) and in the presence of 1 mM ZMP (bottom). Intensities of Cy3 (green), Cy5 (red), and the resulting *E*_FRET_ (blue) are shown. (**c**) The percentage of ZMP-bound aptamer plotted against ZMP concentration and fitted to the Langmuir function for thermally refolded (black) and vectorially folded (red) Δterm. The apparent *K*_D_ and maximum percentage (*P*_max_) were determined to be 1.3 μM and 71%, respectively, for thermally refolded and 1.5 μM and 73% for vectorially folded. (**d**) *E*_FRET_ histograms of Δterm after VF at 0 mM ZMP and incubation with different concentrations of ZMP. Here, a multiple turnover VF protocol was used to maximize the amount of unwound Δterm. Global fits were performed as in b.

Consistent with equilibrium binding measurements (Fig. S1b) in which ZMP is unable to bind to 3’ extended ZTP riboswitch RNAs, the 94-nt wild-type ZTP riboswitch (WT) containing the terminator sequence yielded predominantly a ZMP-unresponsive “terminated” conformation (*E*_FRET_ ~0.2, Fig. S2a) after being thermally refolded in the absence of ZMP. Thermal refolding of the WT construct in the presence of 1 mM ZMP yielded only ~10% ZMP-bound aptamer-containing conformation (*E*_FRET_ ~0.5, Fig. S2a). These data suggest that the aptamer-containing conformation cannot be efficiently accessed when the fully synthesized riboswitch is refolded after thermal denaturation either in the absence of presence of ZMP.

To compare with thermal refolding, we folded the aptamer-only Δterm and terminator-containing WT construct using a helicase-based vectorial folding assay (Fig. 2d, 3a and Online Methods).(Hua et al., 2018a) The assay mimics cotranscriptional RNA folding by unwinding an RNA/DNA heteroduplex with the extremely processive engineered DNA helicase Rep-X(Arslan et al., 2015), which releases the RNA strand sequentially in the 5’-to-3’ direction, the direction of RNA synthesis during transcription (Fig. 1b). An ~0 *E*_FRET_ state was observed before initiating unwinding by ATP addition (Fig. 3a), consistent with separation of the fluorophores by ~60 base pairs (~20 nm). When the WT RNA was vectorially folded in the presence of ≤ 0.01 mM ZMP, only the low-*E*_FRET_ (~0.2) terminated conformation was observed. At higher ZMP concentrations, we observed the mid-*E*_FRET_ (~0.5) aptamer-containing conformation. At 0.1 mM and 1 mM ZMP, 17% and 38%, respectively, of all released RNA showed the aptamer-containing conformation (Fig. 3a), higher than the percentage obtained by refolding of the WT construct (Fig. S2a). These results are consistent with bulk single-round transcription termination assays where riboswitch activation during transcription requires at least 0.2 mM ZMP, a concentration far higher than the *K*_d_ of ~0.5 μM for the isolated aptamer at equilibrium (Fig. 3b). In contrast, different folding approaches did not affect the folding outcome of the aptamer-alone Δterm construct, as the vectorially folded and thermally refolded Δterm bound ZMP to a similar affinity and extent (Fig. 2c). The ZMP-bound aptamer-containing conformation remained for at least 10 min after unwinding, suggesting that the conformation is stable during our measurements (Fig. S2b,c). In the absence of ZMP, transition from the ZMP-unbound aptamer-containing conformation into the terminated conformation must occur rapidly, as addition of 1 mM ZMP 30 s after initiating unwinding did not reveal any ligand-responsive subpopulation (Fig. S2d).

**Figure 3.**
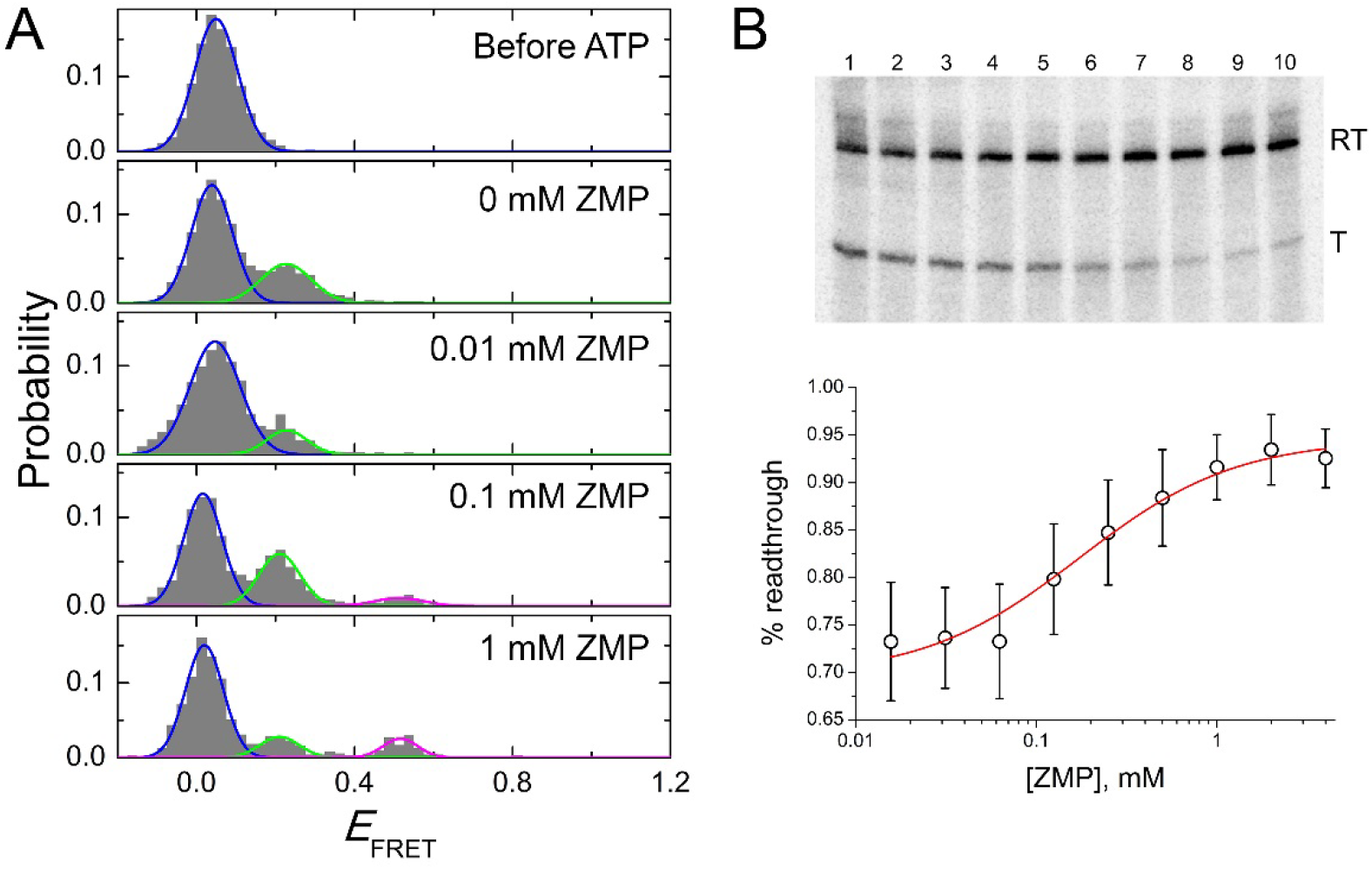
Single-molecule helicase-catalyzed unwinding experiments of the *F. ulcerans* ZTP riboswitch. (**a**) The *E*_FRET_ histograms of the WT heteroduplex before and after ATP addition at different ZMP concentrations using VF with Rep-X. Gaussian fitting with global constraints was used to determine the relative populations of the heteroduplex (blue), terminator (green) and ZMP-bound aptamer (magenta) conformations. (**b**) Bulk single-round transcription termination experiments of ZTP riboswitch are shown in the presence of 0.2 mM NTPs and varying amounts of ZMP. Quantification of all experiments under these conditions is shown below (*mean* ± *standard deviation* (*s.d.*), *n* = 6 independent experiments).

### Effects of unwinding velocity, pausing, and terminator stability on riboswitch folding

To test the effect of unwinding speed, which can be used to mimic transcription speed, on riboswitch folding, we replaced Rep-X with the helicase PcrA-X, which was previously shown to unwind dsDNA substrates at an ~10-fold slower rate.(Arslan et al., 2015) Compared to Rep-X, PcrA-X yielded a higher fraction of the WT construct adopting the mid-*E*_FRET_ aptamer folded state, especially at 0.01-0.1 mM ZMP (Fig. 4a,b). In bulk transcription assays, the ZMP concentration required for riboswitch activation also decreased as the apparent rate of transcription decreased at lower NTP concentrations (Fig. S2e,f,g). Overall, less ZMP is required for riboswitch activation when more time is allowed for ligand binding, consistent with a kinetic control model for ZTP riboswitch function under these conditions.

**Figure 4.**
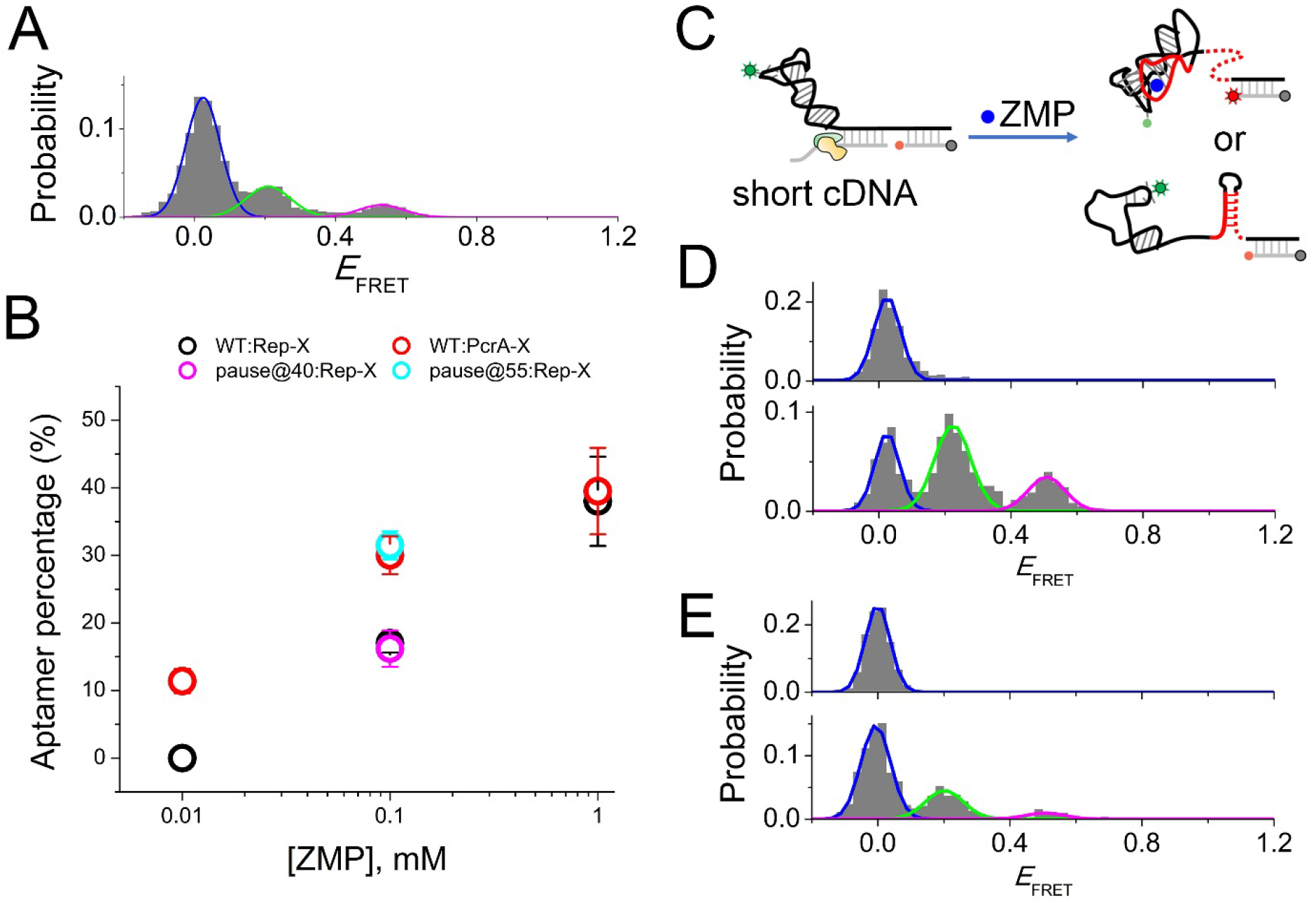
The effect of slower unwinding speed and artificial pausing on riboswitch folding. (**a**) VF histograms for WT ZTP riboswitch unwound by PcrA-X in the presence of 0.1 mM ZMP are shown with Gaussian fits to heteroduplex (blue), terminator (green), and aptamer (magenta) populations. (**b**) The percentage of aptamer-containing conformation obtained by VF with Rep-X (black) and PcrA-X (red) are shown in the presence of 0.01 to 1 mM ZMP (*mean* ± *s.e.m.*, *n* = 3-4). The percentage of aptamer-containing conformation obtained by VF for pause mimics at positions 40 and 55 (magenta and cyan, respectively) are shown (*mean* ± *s.e.m.*, *n* = 2-4). (**c**) Cartoon depicting the pause mimic complex at position 54, in which a cDNA is hybridized only to nt 55-94 of the WT ZTP riboswitch. (**d**) *E*_FRET_ histograms of the cDNA:WT heteroduplexes mimicking a pause at position 55 before (top) and after (bottom) addition of ATP in the presence of 0.1 mM ZMP, as measured by VF with Rep-X. (**e**) *E*_FRET_ histograms of the pause-mimicking heteroduplexes at position 40 before (top) and after (bottom) addition of ATP in the presence of 0.1 mM ZMP, as measured by VF with Rep-X.

Similarly, transcription pausing could provide additional time for aptamer domain folding and ligand binding.(Chauvier et al., 2017; Perdrizet et al., 2012) For example, if cotranscriptional folding of helices P1-2 (nt 1-54) is slow, a pause at nt 54 would promote aptamer formation. In cotranscriptional studies using RNAP,(Frieda and Block, 2012; Uhm et al., 2018) it is difficult to introduce pauses at arbitrary positions. In VF experiments, we mimic the pause at nt 54 by using a DNA that hybridizes only to nt 55-94 of the WT riboswitch, allowing nt 1-54 to pre-fold (Fig. 4c). When this shorter heteroduplex was vectorially folded, an ~2-fold higher percentage of WT adopted the aptamer-containing conformation under limiting ZMP concentration of 0.1 mM, Fig. 4d). This increase in aptamer-containing conformation was similar to what was achieved by VF with the slower PcrA-X (Fig. 4b). In contrast, when we introduced a pause at nt 40, which allows helix P2 but not P1 to equilibrate, the percentage of aptamer-containing conformation remained the same as the full-length heteroduplex (Fig. 4b,e). These results suggest that the slow folding of helix P1 is rate-limiting for aptamer formation. Indeed, sequential folding of the ZTP riboswitch sequence *in silico* predicts a non-native 6-bp helix to stay folded until nt 54, at which point the 6-bp helix is predict to dissolve in favor of P1 helix folding (Fig. S3a). Recent chemical probing of different lengths of the *Clostridium beijerinckii* ZTP riboswitch directly observed the formation of this helix, which occurs in half of ZTP riboswitches.(Strobel et al., 2019) Natural ZTP riboswitch sequences vary significantly in the sequence after nt 54 and often contain additional helices that may modulate RNAP through pausing.

Next, we tested how terminator stability affected riboswitch function. In bulk transcription assays, mutation of the first three terminator bases (81-83A) reduced the ZMP concentration required for riboswitch activation and led to substantially higher readthrough compared to WT (Fig. S3b,c). Consistently, VF experiments with the 81-83A mutant observed a larger fraction of RNAs in the ZMP-bound aptamer-containing conformation (*E*_FRET_ ~0.4, Fig. S3d,e). The 81-83A mutation did not notably reduce the propensity of fully synthesized RNAs to form the terminator, as 81-83A still yielded predominately the terminated conformation upon thermal refolding (~85% *vs.* ~90% for WT, Fig. S2a and S3f). In contrast, mutation of the last three terminator bases (92-94A) caused more pronounced destabilization of the terminated conformation, as 92-94A was partially responsive to ZMP at equilibrium after thermal refolding (Fig. S3g). Moreover, the double mutant (81-83,92-94A) destabilizes the terminator so significantly that the majority of this mutant binds ZMP (Fig. S3h). Taken together, disruption of different terminator regions affects folding and switch outcomes both by decreasing stability and possibly by kinetic factors, which is further explored below.

### Single-molecule fluorescence measurements of unwinding speed

As the unwinding speed is a critical parameter for relating VF to cotranscriptional folding, and as previous Rep-X rate measurements examined dsDNA templates rather than the RNA/DNA hybrid duplexes, we measured helicase speed in three independent ways (Fig. 5a). First, we extended the DNA strand in the heteroduplex to accommodate an additional 18-nt Cy3-labeled oligo (Fig. 5a, top). Upon initiating unwinding, Rep-X first displaced the 18-nt oligo, resulting in ~50-percent decrease in Cy3 intensity (Fig. 5b). Subsequently, we observed an increase and then a decrease in Cy3 intensity which we attributed to protein-induced fluorescence enhancement (PIFE)(Fischer et al., 2004; Hwang and Myong, 2014) caused by helicase unwinding through the Cy3 labeling position at U32. PIFE peak centers were determined by Gaussian fitting and used to define the position of Rep-X on the template (Online Methods). The time difference (Δt) measured between the Cy3 intensity drop and PIFE peak center corresponds to the time for Rep-X to unwind 32 base pairs of the heteroduplex, from which we estimated the unwinding speed to be 65.0 ± 0.39 nt/s (Fig. 5c).

**Figure 5.**
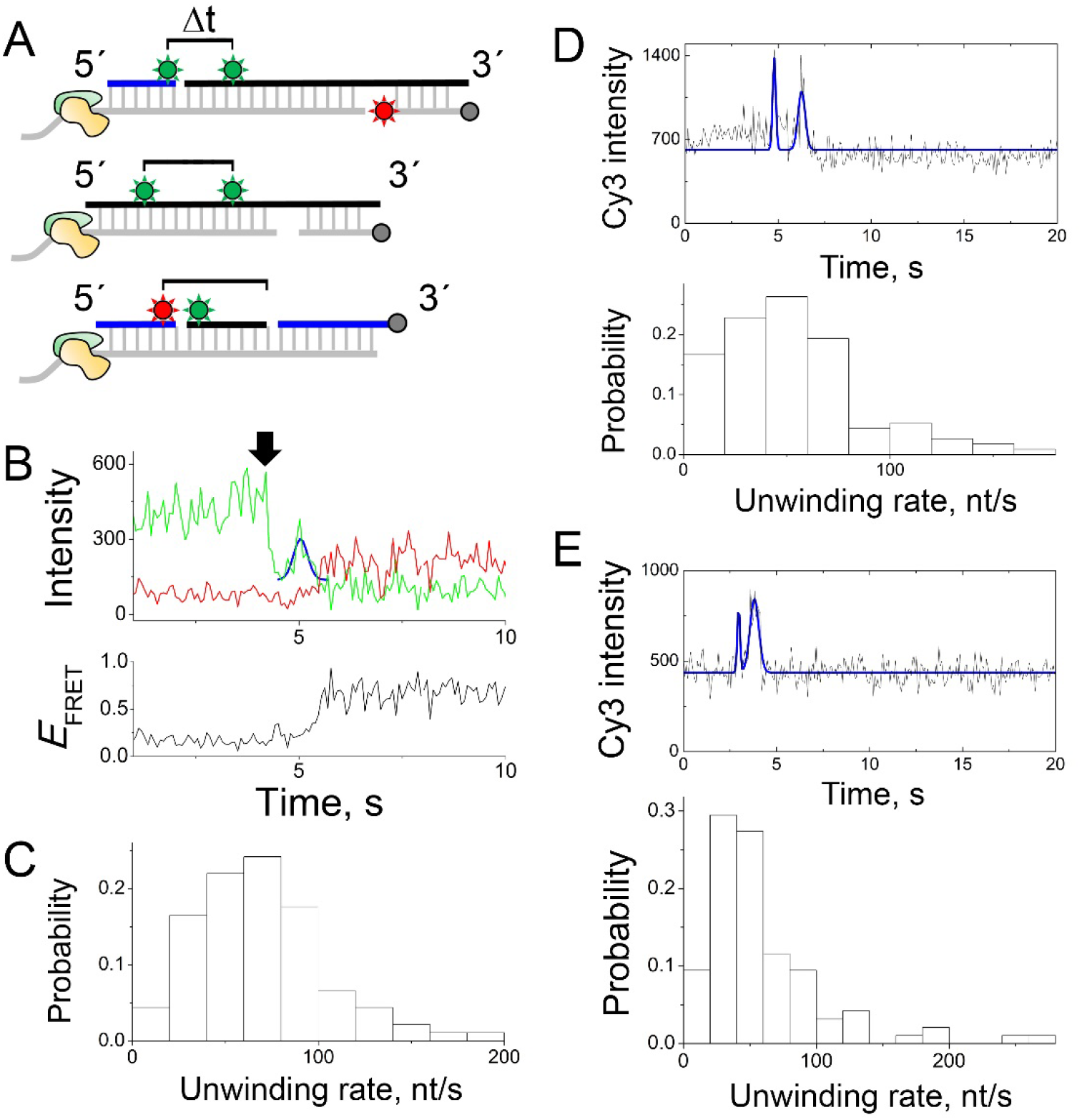
Real-time helicase rate measurements (**a**) Cartoon depicting the three constructs used for rate measurement experiments, either through the loss of Cy3 intensity and PIFE (top), two PIFE peaks (middle), or the loss of FRET and Cy3 intensity (bottom). The cartoon shows Rep-X helicase in light yellow and light green, DNA strands in grey, ZTP riboswitch-derived RNA strands in black, other DNA or RNA strands in blue, Cy3 and Cy5 dyes indicated by green and red stars, and biotin labels indicated by grey circles. A black bracket indicates the two events used to measure helicase unwinding rates. (**b**) Individual single-molecule trajectory in the presence of 0.1 mM ZMP for the top scheme, using loss of Cy3 intensity and PIFE. The drop in Cy3 intensity is arrowed, and the PIFE peak fit by a Gaussian curve (blue line). (**c**) Histogram of helicase rates obtained from all such experiments. The Δt between intensity drop and PIFE peak is shown (*mean* ± *s.e.m.*, *n* = 91 molecules). (**d**) Individual single molecule trajectory (top) and histogram of helicase rates (bottom) for double PIFE experiment with Cy3 labels at U12 and U84. For the example trajectory, the raw data (black) and Gaussian fits of PIFE peaks (blue) are shown. The mean time difference (Δt) between PIFE peaks for experiments is 2.01 ± 0.20 s (*mean* ± *s.e.m.*, *n* = 95 molecules). (**e**) Individual single molecule trajectory (top) and histogram of all helicase rates (bottom) for double PIFE experiment with Cy3 labels at U32 and U84. The Δt between PIFE peaks is 1.64 ± 0.15 s (*mean* ± *s.e.m.*, *n* = 112 molecules).

In a second approach to measure unwinding speed, we positioned two Cy3 fluorophores along the WT riboswitch sequence and excluded the Cy5 fluorophore (Fig. 5a, middle). This labeling strategy resulted in two PIFE peaks during unwinding (Fig. 5d,e), and the Δt between the two peak centers, which corresponds to the time for Rep-X to unwind from the first Cy3 position to the second, allowed estimation of unwinding speed using only a single fluorophore. Separation of the dyes by 72 base pairs (Fig. 5d) and 52 base pairs (Fig. 5e) resulted in average time differences of of 2.01 ± 0.20 s and 1.64 ± 0.15 s, respectively, and unwinding speeds of 60.4 ± 4.8 nt/s and 52.5 ± 3.1 nt/s, respectively.

As the 3’ end of the ZTP riboswitch sequence is more GC-rich than the 5’ end, we used a third method to measure the unwinding speed over different portions of the duplex (Fig. 5a, bottom). In this method, unwinding releases a Cy5-labeled DNA oligo prior to release of the Cy3-labeled RNA oligo, which is observed by a loss in FRET prior to a loss in Cy3 intensity (Fig. S4a,b). The average unwinding speed over different portions of the WT riboswitch sequence (Fig. 5a,bottom) was similar (Fig. S4a), suggesting Rep-X unwinds at a uniform average velocity over the duplex. Importantly, the loss of FRET in this assay is easily distinguished from photobleaching when many traces are examined (Fig. S4c), despite appearing identical for any particular single trajectory in this experiment. Also, the average time interval between loss of the first and second oligos in these experiments was unchanged when an RNA oligo was replaced by a DNA oligo and vice versa (Fig. S4d,e). Compared to Rep-X, PcrA-X facilitated duplex unwinding more slowly, at a speed of ~8 nt/s (Fig. S4b), which was also distinguished from photobleaching when many traces were examined (Fig. S4f). Thus, the ~6-fold change in unwinding speed between PcrA-X and Rep-X is significant enough to modulate the percentage of aptamer-containing conformation obtained after VF.

### Real time unwinding measurements to observe folding pathways

To investigate how folding dynamics affects termination, we performed single-molecule experiments to monitor the decision-making trajectory of individual riboswitches in real time (Online Methods). As in above experiments, we observed an intensity peak in the Cy3 signal, which we attributed to PIFE(Hwang and Myong, 2014) upon helicase unwinding through the Cy3 labeling position at U32 (Fig. 6, S5). The center of each PIFE peak was determined by Gaussian fitting, and trajectories were synchronized and aligned according to PIFE peak centers, which is critical for comparing trajectories. In the presence of 1 mM ZMP, WT and 81-83A molecules folded into and remained for long periods in the higher *E*_FRET_ aptamer-containing conformation (Fig. 6a and S5a). In the absence of ZMP, most WT molecules were observed to transition into the low-*E*_FRET_ (~0.2) terminated conformation within 1.5 s; however, 8% (2/24) of the WT trajectories showed high-*E*_FRET_ values (≥ 0.6) for ≥ 3 s (Fig. 6b), longer than what it takes for Rep-X to complete unwinding given its speed. Although 81-83A predominantly formed the terminated conformation 1 min after initiating unwinding (Fig. S6c), 60% (9/15) of the 81-83A trajectories maintained these high-*E*_FRET_ conformations for ≥ 3 s in the initial phase of folding (Fig. 6c), suggesting formation of the mutant terminator was severely delayed during VF. Such events, which could functionally promote transcription readthrough, provide a plausible mechanism for “leaky” readthrough (*i.e.*, inefficient termination in the absence of ligand) and explain how the 81-83A mutation affects termination efficiency in bulk transcription assays despite folding primarily into a terminated state under equilibrium refolding measurements.

**Figure 6.**
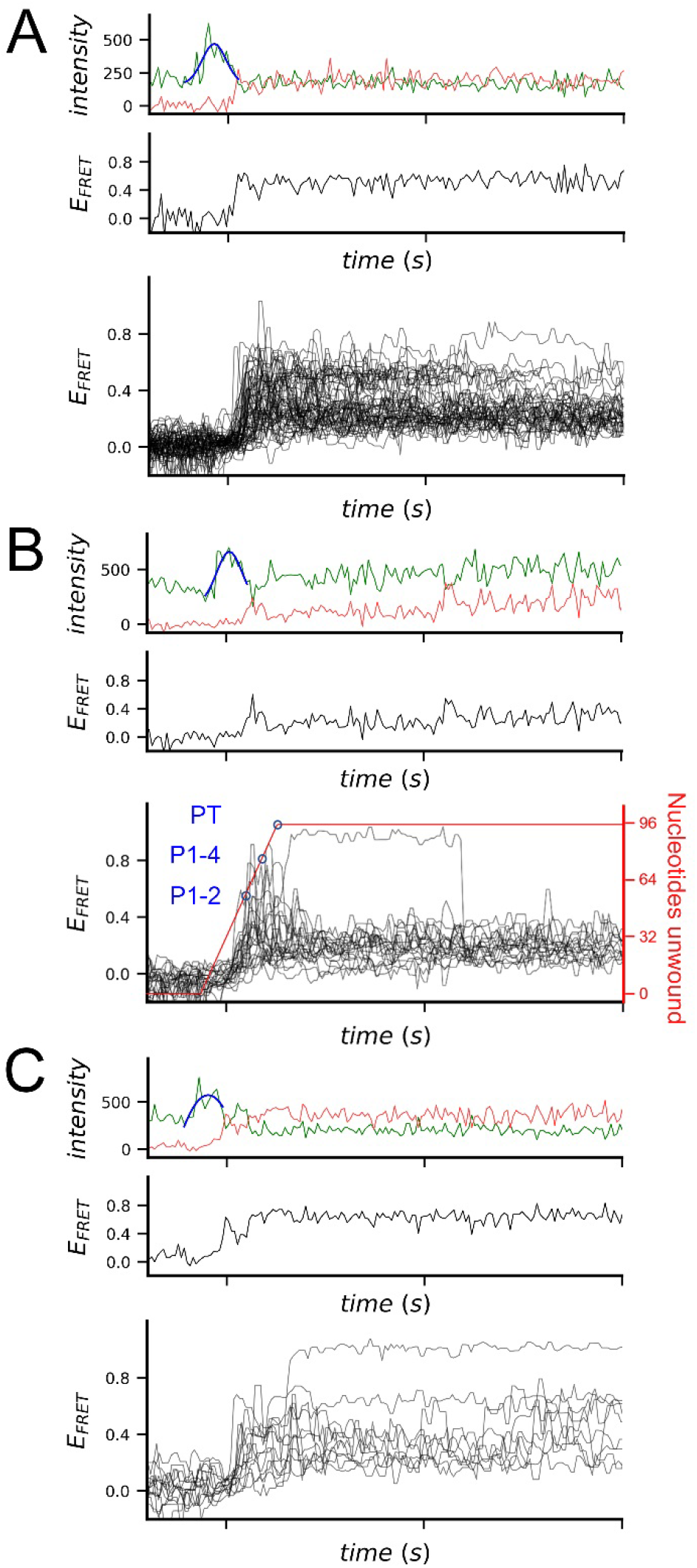
Real-time observation of heteroduplex unwinding and riboswitch folding in VF assays. (**a**) Individual real-time trajectory (top) and overlay of all WT ZTP riboswitch folding trajectories (bottom, *n* = 56) using VF with Rep-X in the presence of 1 mM ZMP. The PIFE peaks were fit to a Gaussian (blue) to determine their centers for synchronizing trajectories. (**b**) Individual real-time trajectory (top) and overlay of all WT ZTP riboswitch trajectories (bottom, *n* = 24) using VF with Rep-X in the absence of ZMP. The relative unwinding position is indicated by a red line, based on the experimentally determined Rep-X unwinding rate (Fig. 3) and the location of U32 on the template. Blue circles indicate time points corresponding to the unwinding and release of helices P1 and P2, helices P1-P4, and terminator helix PT. (**c**) Real-time trajectories of terminator mutant 81-83A in the absence of ZMP, showing an individual trajectory (top) and all trajectories overlaid (bottom, *n* = 15). The time duration between ticks is 5 s.

## Discussion

From these experiments, an individual riboswitch is observed to fold through one of many distinct channels, and the entire folding landscape is made evident from many observations (Fig. 6). Even delayed folding events, like those identified here, may increase leaky readthrough for the purpose of tuning the landscape to a desired threshold with higher basal activation, such that the gene is partially ON even in the absence of ligand. In both bulk transcription and single-molecule unwinding assays, only ~40% of this riboswitch is modulated by ligand binding, suggesting that a large fraction of RNA falls into leaky terminated or leaky readthrough states. In bulk transcription assays, “leaky” readthrough decreases as transcription speed slows at lower NTP concentrations (Fig. S2e,g). What is not evident from bulk assays, however, is how alternative folding plays a role in populating leaky states.

In order to relate RNA folding during helicase unwinding experiments to RNA folding during transcription, the rates of unwinding and transcription should be similar. The three approaches we have used to investigate unwinding speed (Fig. 5) show that Rep-X releases the RNA at a rate of ~60 nt/s, which is ~2.5-fold slower than the previously reported rate for Rep-X on dsDNA substrates(Arslan et al., 2015) but within the 20-80 nt/s synthesis rate for bacterial RNA polymerases.(Pan and Sosnick, 2006) The advantages of the first method to determine the speed (Fig. 5b,c) are that unwinding speed and RNA folding can be measured in the same experiment to control for varying experimental conditions that might affect helicase speed. The drop in Cy3 intensity followed by a PIFE signal is an easily identified mark at the start of the trajectory. In comparison, using multiple PIFE signals to assay unwinding speed (Fig. 5d,e) utilizes substrates that are easily prepared and measured via only Cy3 excitation but do not report on RNA folding. This method is advantageous when only the speed is of interest and could be used for any enzyme moving in a single direction along the substrate, provided that PIFE is observed.

Under these conditions, Rep-X is well suited for mimicking bacterial RNAP transcription speeds. While the isolated aptamer domain displays reversible binding behavior and could be just as well investigated by equilibrium folding measurements (Fig. 2), the terminator-containing WT construct must be assayed during unwinding to investigate the switching behavior (Fig. 3), as it is kinetically controlled in both bulk transcription assays and single-molecule unwinding assays. Consequently, pausing and slower unwinding speed both promote ligand-responsive switching as both allow the RNA longer time to fold before the terminator element is produced. As kinetically controlled switching behavior is observed for this riboswitch with two different motile enzymes (Rep-X and RNAP), the RNA sequence itself contains the kinetic control information, which is interpreted by the motile enzyme. Consequently, a conserved riboswitch sequence could still function kinetically in the context of different cellular processes (*i.e.*, transcription, translation), as long as a motile enzyme is present to impose sequential movement.

In summary, we applied a novel superhelicase-based VF assay to study the *F. ulcerans* ZTP riboswitch, whose folding is similarly rate-sensitive during both transcription and helicase-catalyzed unwinding. Kinetically controlled ligand-responsive switching is observed as long as a motile enzyme (either RNAP or helicase) sequentially releases the functional RNA, which would otherwise fold into a stable terminated conformation incapable of switching. As these motile enzymes function at similar rates (~60 nt/s), the VF assay could be widely adapted to study other cotranscriptional cellular processes such as assembly of ribonucleoprotein complexes and even telomere folding and assembly.(Mitra and Ha, 2019b)

## STAR METHODS

Detailed methods are provided in the online version of this paper and include the following:

- KEY RESOURCES TABLE
- CONTACT FOR REAGENT AND RESOURCE SHARING
- EXPERIMENTAL MODEL AND SUBJECT DETAILS
- METHOD DETAILS
  - Sample preparation for single molecule measurements
  - Single molecule measurements
  - Single-molecule data acquisition and analysis
  - Single-round transcription termination assays
  - Isothermal calorimetry
  - RNA and DNA oligonucleotide sequences
- QUANTIFICATION AND STATISTICAL ANALYSIS

## SUPPLEMENTAL INFORMATION

Supplemental Information contains five figures and one table and can be found online with this article at [link].

## ACKNOWLEDGMENTS

We thank Drs. G. Piszczek and D. Wu in the NHLBI Biophysics Core for ITC support. This work was supported by U.S. NSF grant PHY 1430124 and NIH grant GM 112659 to T.H. and NIH grant K22HL139920-01 to C.P.J. T.H. is an investigator of the Howard Hughes Medical Institute. This work was supported in part by the intramural program of the NHLBI, NIH.

## AUTHOR CONTRIBUTIONS

B.H., C.P.J., A.R.F. and T.H. designed the project, B.H., J.M., P.J.M, and R.R. performed the single-molecule experiments, C.P.J. prepared single-molecule constructs and performed bulk experiments, B.H. and C.P.J. analyzed the data, and B.H., C.P.J., A.R.F. and T.H. wrote the manuscript.

## STAR METHODS

### KEY RESOURCES TABLE

| REAGENT or RESOURCE | SOURCE | IDENTIFIER |
| --- | --- | --- |
| **Chemicals, Peptides, and Recombinant Proteins** |  |  |
| Rep-X helicase | Arslan et al, *Science* (2015) | N/A |
| PcrA-X helicase | Arslan et al, *Science* (2015) | N/A |
| T4 RNA ligase 2 | New England Biolabs | Cat# M0239L |
| E. coli RNA polymerase, holoenzyme | New England Biolabs | Cat# M0551S |
| Trolox | Sigma-Aldrich | Cat# 238813 |
| Glucose oxidase | Sigma-Aldrich | Cat# G2133 |
| Catalase | Sigma-Aldrich | Cat# C1345 |
| Neutravidin | Thermo Scientific | Cat# 31000 |
| **Oligonucleotides** |  |  |
| See Table S1 for all RNA and DNA oligonucleotides. |  |  |
| **Recombinant DNA** |  |  |
| See Table S1 for partial plasmid sequences. |  |  |
| *F. ulcerans* ZTP aptamer plasmid | Jones and Ferré-D'Amaré, *NSMB* 2015 | N/A |
| *F. ulcerans* ZTP transcription termination plasmid | This paper | N/A |
| *F. ulcerans* ZTP transcription termination plasmid G81A | This paper | N/A |
| *F. ulcerans* ZTP transcription termination plasmid 81-83A | This paper | N/A |
| *F. ulcerans* ZTP transcription termination plasmid 81-86A,87C | This paper | N/A |
| **Software and Algorithms** |  |  |
| Origin 8 | OriginLab Corp. | https://www.originlab.com/ |
| Single-molecule fluorescence acquisition software | Custom made | Available from the Ha lab |
| MATLAB | The MathWorks, Inc. | https://www.mathworks.com/ |

#### Sample preparation for single molecule measurements

A detailed description of preparation of fluorescently labeled RNA and RNA-DNA complexes for has been described (Jones et al., 2019a). All steps involving fluorescently labeled oligomers were performed in the dark. Unlabeled DNA oligomers, primers and plasmids (Supplementary Table 1) were purchased from Eurofins Genomics LLC and Integrated DNA Technologies. Fluorescently labeled oligonucleotides were purchased from Trilink (San Diego, CA), including the Cy3-labeled 5’-end segment of the *F. ulcerans* ZTP riboswitch (nt 1–33, Supplementary Table 1) and the Cy5-labeled and biotinylated tether DNA (biotin-DNA-Cy5, Supplementary Table 1). Fluorophores were attached to these oligomers via 6-carbon amino linkers at the C5 position of U32 and at the 3’-end of the biotinylated DNA, respectively. ZTP riboswitch RNAs lacking the first 33 nucleotides (Δ1–33) were transcribed and purified from DNA templates PCR amplified from plasmids, as previously described (Ferré-D’Amaré and Doudna, 1996; Jones and Ferré-D’Amaré, 2015). Mutations to the terminator stem were made via PCR amplification using a reverse primer containing the mutations. Δ1–33 RNAs were phosphorylated with T4 polynucleotide kinase (New England Biolabs), phenol-chloroform extracted, and ethanol precipitated. The Cy3-labeled nt 1–33 RNA and Δ1–33 RNAs were annealed to a DNA splint identical to the cDNA-dT_20_ used in single molecule measurements and ligated using T4 RNA ligase 2 (New England Biolabs). Ligated RNAs were purified by 8% denaturing urea-PAGE, recovered via elution in Whatman Elutrap electroelution systems, concentrated, washed with DEPC-treater water, and filtered in 0.22 μm spin filters. Concentrations were calculated by the absorbance of Cy3 at 552 nm (ε_552_ = 0.15 μM^−1^ cm^−1^) and the absorbance of the RNAs at 260 nm (ε_260_ = 1.16 μM^−1^ cm^−1^ for the terminator-containing RNAs and ε_260_ = 0.982 μM^−1^ cm^−1^ for the aptamer-only RNA) after correcting for the contribution of Cy3 absorbance at 260 nm (~5 % correction). Ligated RNAs were aliquoted and stored at −20 °C in the dark until further use.

Fluorescently labeled RNAs for double PIFE helicase rate measurements (Fig. 3d, S6) were constructed in a similar manner, resulting in RNAs identical to the WT sequence and containing two Cy3 labels. For these experiments, three RNAs were ligated, consisting of segments spanning riboswitch nucleotides 1-33, 34-71, and 72-94. The segment 1-33 contained a Cy3 label either at position U12 or position U32. The segment 72-94 contained a Cy3 label at position U84 and also contained the RNA sequence complementary to the biotinylated DNA.

To form annealed heteroduplexes for single-molecule measurements, 20 μl reaction aliquots containing a 1:0.8:10 ratio of Cy3-RNA:biotin-DNA-Cy5:cDNA-dT_20_ (160 nM Cy3-RNA, 128 nM biotin-DNA-Cy5, and 1.6 μM cDNA-dT_20_) were prepared in the buffer containing 50 mM HEPES-KOH (pH 7.4) and 150 mM KCl. The reaction aliquots were slowly cooled from 95 °C to 20 °C in a thermocycler (30 s/5 °C). Annealing success was evaluated on 8% nondenaturing PAGE (in 1X TBE) by comparison to reactions lacking either biotin-DNA-Cy5, cDNA-dT_20_, or both. Reactions lacking cDNA-dT_20_ were also saved for single molecule measurements. Gels were scanned with a Typhoon Trio Variable Mode Imager (GE Healthcare) to identify bands containing fluorescent labels.

For dropoff-PIFE experiments (Fig. 4b,c), a longer cDNA-dT_20_ (cDNA-speedmer-dT_20_) was used for annealing reactions in the presence of an 18-nt RNA oligo labeled at the 3’ end with Cy3 (18-speedmer), the WT Cy3-labeled RNA, and biotin-DNA-Cy5. Reactions contained a 1:0.8:9:10 ratio of Cy3-RNA:biotin-DNA-Cy5:cDNA-speedmer-dT_20_:18-speedmer to saturate cDNA-speedmer-dT_20_ with 18-speedmer. Annealing success was evaluated via nondenaturing PAGE as above.

For dropoff experiments (Fig. S7), 20 μl reaction aliquots containing a 1:1.2:1.21.2 ratio of biotin-DNA:cDNA-dT_20_:Cy3-oligo:Cy5-oligo (1 μM biotin-DNA, 1.2 μM cDNA-dT_20_, 1.2 μM Cy3-oligo and 1.2 μM Cy5-oligo) were prepared in the buffer containing 50 mM HEPES-KOH (pH 7.4) and 150 mM KCl. The reaction aliquots were slowly cooled from 95 °C to 20 °C in a thermocycler (30 s/5 °C). By selecting surface-tethered molecules that are doubly labeled, we ensured the molecules being imaged contain all four strands.

#### Single molecule measurements

Vectorial folding (VF) was performed as previously described (Hua et al., 2018a) with the following modifications. Rep-X was incubated in the loading buffer for 5 min with heteroduplexes that were immobilized on the imaging surface. The loading buffer contained 50 nM Rep-X, 50 mM HEPES-KOH (pH 7.4), 150 mM KCl and 10 mM MgCl_2_. Free Rep-X was washed out and unwinding was initiated by adding the unwinding buffer. The unwinding buffer contained 50 mM HEPES (pH 7.4), 150 mM KCl, 10 mM MgCl_2_, 1 mM ATP and different concentrations of ZMP. To stabilize the remaining aptamer conformation and distinguish it from the terminator conformation, 1 mM ZMP was supplied into the imaging channel after 1 min of vectorial folding. To observe the heteroduplex unwinding and riboswitch folding in real-time, imaging was started ~5 s before the addition of the unwinding buffer. The loading and unwinding buffers used during imaging contained additional 4 mM Trolox, 0.8 % wt/vol glucose, 165 U/ml glucose oxidase and 2170 U/ml catalase to reduce the photobleaching rate (Rasnik et al., 2006).

For VF multiple turnover experiments with Rep-X (Fig. 2d), 50 nM Rep-X was included in the unwinding buffer and no separate Rep-X loading step was performed. When PcrA-X was used in place of Rep-X, 100 nM of PcrA-X was added in the loading buffer. Other buffer components were kept the same.

To refold RNAs for single-molecule measurements, 20 μl reaction aliquots containing a 1:0.8 ratio of Cy3-RNA:biotin-DNA-Cy5 (160 nM Cy3-RNA and 128 nM biotin-DNA-Cy5) were prepared in the buffer containing 50 mM HEPES-KOH (pH 7.4), 150 mM KCl, 10 mM MgCl_2_ and different concentrations of ZMP. The reaction aliquots were slowly cooled from 95 °C to 20 °C in a thermocycler (30 s per 5 °C step). When refolded at 1 mM ZMP, RNAs were imaged within 5 min after refolding to minimize the loss of ZMP-bound aptamer due to terminator hairpin formation.

#### Single-molecule data acquisition and analysis

Single-molecule data acquisition and analysis were performed as previously described (Hua et al., 2018a; Hua et al., 2018b). To plot the trajectory overlay, we synchronized trajectories according to the PIFE peak center. The percentage of aptamer-like fold is calculated as the ratio of the aptamer population (magenta) over the sum of the terminator (green) and aptamer populations.

For traces containing multiple PIFE peaks (Fig. 3d, S6), peak centers were determined by Gaussian fitting in Origin (OriginLab, Northampton, MA), and time differences between PIFEs (Δt) were calculated by the difference between the peak centers. Traces appearing to contain too many PIFE peak were excluded from analysis (7 traces excluded from 12-84 dataset, 8 traces from 32-84 dataset, and 9 traces from drop-PIFE dataset). Unwinding rates are reported as the *mean* ± *s.e.m*. with the *n* being the number of individual molecules.

For dropoff-PIFE traces (Fig. 4b,c), the drop in Cy3 intensity was determined by eye, and time was recorded. PIFE peak centers were determined by Gaussian fitting as above. The time difference (Δt) between these two events was calculated for each trace and used to determine a helicase unwining rate (in nt/s) with the known distance of 34 nt separating the two events. Traces containing more than one PIFE peak were excluded from analysis (9 traces excluded). The average rate is reported as the *mean* ± *s.e.m.* with *n* being the number of individual molecules (91 total).

For dropoff experiments (Fig. S7), the decrease in FRET value and decrease in Cy3 intensity were determined by eye. The time difference (Δt) between these two events was calculated for each trace, and the average unwinding rate was calculated using all traces for which Δt was less than 2 s for Rep-X (or 10 s for PcrA-X), which includes traces contained within the Gaussian distribution of rates.

#### Single-round transcription termination assays

PCR templates were amplified from a plasmid containing a 100 nt leader sequence, a λ Pr promoter, 26-nt C-less region, the *F. ulcerans* ZTP riboswitch sequence, which contains the aptamer domain, expression platform and 51 nucleotides beyond the terminator U8 sequence. After amplification, templates were purified by 2% agarose gel electrophoresis. Mutations to the templates were made by site-directed mutagenesis to the plasmid and confirmed by sequencing.

Transcription reactions were performed by halting transcription by omission of CTP as previously described and restarting transcription by adding NTPs and ZMP (Jones and Ferré-D’Amaré, 2015). Halted transcription complexes contained 80 pmol uL^−1^ DNA template, 20 mM Tris-HCl, pH 8, 20 mM NaCl, 4 mM MgCl_2_, 0.1 mM DTT, 0.1 mM EDTA, 4% glycerol, 0.14 mM ApU, 1 μM GTP, 2.5 μM ATP, 2.5 μM UTP, ~1 μCi mL^−1^ [α-^32^P]-ATP, 0.04 U uL^−1^ *Escherichia coli* RNA polymerase holoenzyme (Epicenter), and 10 μM of a 26 nt oligonucleotide complementary to the C-less region. Transcriptions were restarted by addition of various concentrations of NTPs and ZMP prior to incubation at 37 °C for the duration of the time course, removing aliquots as needed and quenching reactions by addition of loading dye containing 8 M Urea, 20% sucrose, 0.1% SDS, 0.01% bromophenol blue, 0.01% xylene cyanol and placing them on ice or storing them at −20 °C. Reactions were then separated via 8% denaturing urea-PAGE and analyzed as previously described (Jones and Ferré-D’Amaré, 2014; Xiao et al., 2008). To determine apparent reaction rates for appearance of terminated and readthrough transcription products, time courses were fit by single exponential fits using Origin. For single time point titrations, reactions containing different concentrations of ZMP were incubated for 20 min at 37 °C, and results were fit to a 1:1 binding isotherm to determine T50 values.

Sequential folding *in silico* was performed by using Mfold (Zuker, 2003) with different lengths of ZTP riboswitch transcripts as inputs.

#### Isothermal calorimetry

All isothermal calorimetry measurements were performed and analyzed as previously described (Jones et al., 2019b), using a MicroCal iTC200 (Malvern, Egham, UK). Briefly, RNAs were refolded by heating at 95 °C for 2 min and immediately placed on ice. MgCl_2_ was added to 10 mM, the sample was brought up to volume, and the RNA was incubated at 37 °C prior to performing titrations at 37 °C in a buffer containing 50 mM Hepes-KOH, pH 7.4, 150 mM KCl, and 10 mM MgCl_2_. Typically the cell containing 20 μM RNA was titrated with 200 μM ZMP, but higher concentrations were used for weaker binders.

## REFERENCES

Arslan, S., Khafizov, R., Thomas, C.D., Chemla, Y.R., and Ha, T. (2015). Protein structure. Engineering of a superhelicase through conformational control. Science 348, 344–347.

Bochner, B.R., and Ames, B.N. (1982). ZTP (5-amino 4-imidazole carboxamide riboside 5’-triphosphate): a proposed alarmone for 10-formyl-tetrahydrofolate deficiency. Cell 29, 929–937.

Chauvier, A., Picard-Jean, F., Berger-Dancause, J.C., Bastet, L., Naghdi, M.R., Dube, A., Turcotte, P., Perreault, J., and Lafontaine, D.A. (2017). Transcriptional pausing at the translation start site operates as a critical checkpoint for riboswitch regulation. Nature communications 8, 13892.

Ferré-D’Amaré, A.R., and Doudna, J.A. (1996). Use of cis- and trans-ribozymes to remove 5’ and 3’ heterogeneities from milligrams of in vitro transcribed RNA. Nucleic acids research 24, 977–978.

Fischer, C.J., Maluf, N.K., and Lohman, T.M. (2004). Mechanism of ATP-dependent translocation of E.coli UvrD monomers along single-stranded DNA. Journal of molecular biology 344, 1287–1309.

Frieda, K.L., and Block, S.M. (2012). Direct observation of cotranscriptional folding in an adenine riboswitch. Science 338, 397–400.

Garst, A.D., Edwards, A.L., and Batey, R.T. (2011). Riboswitches: structures and mechanisms. Cold Spring Harbor perspectives in biology 3.

Helmling, C., Wacker, A., Wolfinger, M.T., Hofacker, I.L., Hengesbach, M., Fuertig, B., and Schwalbe, H. (2017). NMR structural profiling of transcriptional intermediates reveals riboswitch regulation by metastable RNA conformations. Journal of the American Chemical Society.

Howe, J.A., Wang, H., Fischmann, T.O., Balibar, C.J., Xiao, L., Galgoci, A.M., Malinverni, J.C., Mayhood, T., Villafania, A., Nahvi, A., et al. (2015). Selective small-molecule inhibition of an RNA structural element. Nature 526, 672–677.

Hua, B., Panja, S., Wang, Y., Woodson, S.A., and Ha, T. (2018a). Mimicking Co-Transcriptional RNA Folding Using a Superhelicase. Journal of the American Chemical Society 140, 10067–10070.

Hua, B., Wang, Y., Park, S., Han, K.Y., Singh, D., Kim, J.H., Cheng, W., and Ha, T. (2018b). The Single-Molecule Centroid Localization Algorithm Improves the Accuracy of Fluorescence Binding Assays. Biochemistry 57, 1572–1576.

Hwang, H., and Myong, S. (2014). Protein induced fluorescence enhancement (PIFE) for probing protein-nucleic acid interactions. Chemical Society reviews 43, 1221–1229.

Jones, C.P., and Ferré-D’Amaré, A.R. (2014). Crystal structure of a c-di-AMP riboswitch reveals an internally pseudo-dimeric RNA. The EMBO journal.

Jones, C.P., and Ferré-D’Amaré, A.R. (2015). Recognition of the bacterial alarmone ZMP through long-distance association of two RNA subdomains. Nature structural & molecular biology 22, 679–685.

Jones, C.P., and Ferré-D’Amaré, A.R. (2017). Long-Range Interactions in Riboswitch Control of Gene Expression. Annual review of biophysics.

Jones, C.P., Panja, S., Woodson, S.A., and Ferre-D’Amare, A.R. (2019a). Monitoring co-transcriptional folding of riboswitches through helicase unwinding. Methods in enzymology 623, 209–227.

Jones, C.P., Piszczek, G., and Ferre-D’Amare, A.R. (2019b). Isothermal Titration Calorimetry Measurements of Riboswitch-Ligand Interactions. Methods in molecular biology 1964, 75–87.

Kim, P.B., Nelson, J.W., and Breaker, R.R. (2015). An ancient riboswitch class in bacteria regulates purine biosynthesis and one-carbon metabolism. Molecular cell 57, 317–328.

Mitra, J., and Ha, T. (2019a). Nanomechanics and co-transcriptional folding of Spinach and Mango. Nature communications 10, 4318.

Mitra, J., and Ha, T. (2019b). Streamlining effects of extra telomeric repeat on telomeric DNA folding revealed by fluorescence-force spectroscopy. Nucleic acids research.

Pan, T., and Sosnick, T. (2006). RNA folding during transcription. Annual review of biophysics and biomolecular structure 35, 161–175.

Perdrizet, G.A., 2nd, Artsimovitch, I., Furman, R., Sosnick, T.R., and Pan, T. (2012). Transcriptional pausing coordinates folding of the aptamer domain and the expression platform of a riboswitch. Proceedings of the National Academy of Sciences of the United States of America 109, 3323–3328.

Rasnik, I., McKinney, S.A., and Ha, T. (2006). Nonblinking and long-lasting single-molecule fluorescence imaging. Nature methods 3, 891–893.

Rohlman, C.E., and Matthews, R.G. (1990). Role of purine biosynthetic intermediates in response to folate stress in Escherichia coli. Journal of bacteriology 172, 7200–7210.

Serganov, A., and Nudler, E. (2013). A decade of riboswitches. Cell 152, 17–24.

Strobel, E.J., Cheng, L., Berman, K.E., Carlson, P.D., and Lucks, J.B. (2019). A ligand-gated strand displacement mechanism for ZTP riboswitch transcription control. Nature chemical biology 15, 1067–1076.

Uhm, H., Kang, W., Ha, K.S., Kang, C., and Hohng, S. (2018). Single-molecule FRET studies on the cotranscriptional folding of a thiamine pyrophosphate riboswitch. Proceedings of the National Academy of Sciences of the United States of America 115, 331–336.

Watters, K.E., Strobel, E.J., Yu, A.M., Lis, J.T., and Lucks, J.B. (2016). Cotranscriptional folding of a riboswitch at nucleotide resolution. Nature structural & molecular biology 23, 1124–1131.

Wickiser, J.K., Cheah, M.T., Breaker, R.R., and Crothers, D.M. (2005a). The kinetics of ligand binding by an adenine-sensing riboswitch. Biochemistry 44, 13404–13414.

Wickiser, J.K., Winkler, W.C., Breaker, R.R., and Crothers, D.M. (2005b). The speed of RNA transcription and metabolite binding kinetics operate an FMN riboswitch. Molecular cell 18, 49–60.

Xiao, H., Edwards, T.E., and Ferré-D’Amaré, A.R. (2008). Structural basis for specific, high-affinity tetracycline binding by an in vitro evolved aptamer and artificial riboswitch. Chemistry & biology 15, 1125–1137.

Zhang, J., Lau, M.W., and Ferré-D’Amaré, A.R. (2010). Ribozymes and riboswitches: modulation of RNA function by small molecules. Biochemistry 49, 9123–9131.

Serganov, A., and Nudler, E. (2013). A decade of riboswitches. Cell 152, 17–24.

Zuker, M. (2003). Mfold web server for nucleic acid folding and hybridization prediction. Nucleic acids research 31, 3406–3415.

