## Supplemental File for "Real-time monitoring of cotranscriptional riboswitch folding and switching"

^‡^ Co-first authors

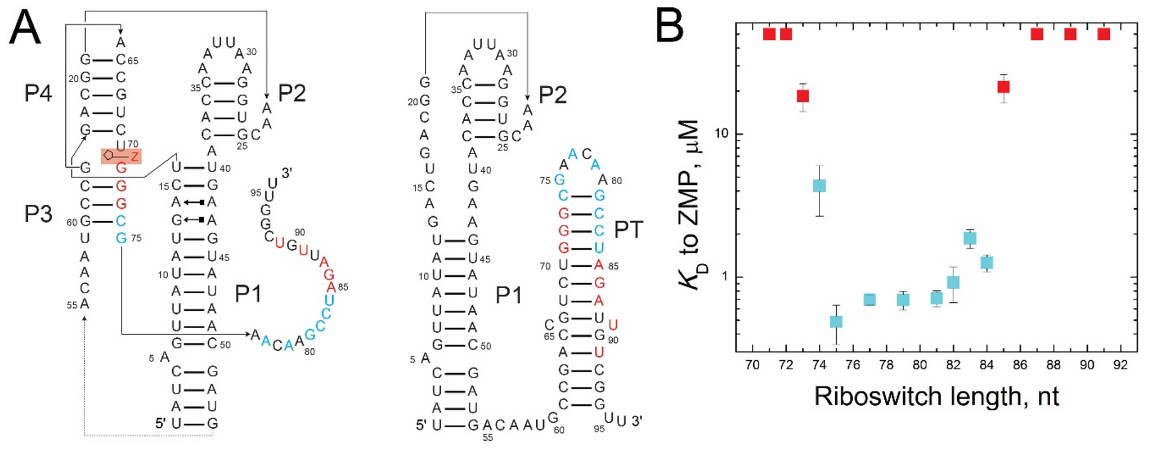

**Supplementary Figure S1 (Related to Figure 2)** ZMP binding to the ZTP riboswitch measured by isothermal titration calorimetry. (**a**) Secondary structures of ZTP riboswitch are shown with the terminator unfolded in the ZMP-bound state (left) and with the terminator (PT) folded in the ZMP-free state (right). Nucleotides are colored to indicate the apparent dissociation constants of the riboswitch variants ending at the respective nucleotides. For example, the 79-nt RNA ending at A79 is colored cyan as this RNA binds ZMP with a *K*_d_ of ~1 μM. (**b**) ZTP riboswitch variants transcribed with different 3´-end lengths were thermally folded and titrated with ZMP to determine apparent dissociation constants via isothermal calorimetry. RNA lengths for which the apparent dissociation constant was tighter than 5 μM are in cyan, and RNA lengths for which the apparent dissociation constant was weaker than 5 μM or for which binding was not observed (i.e., *K*_d_ > 50 μM) are in red. The apparent dissociation constant for the 75-nt RNA is the previously determined value (Jones and Ferré-D’Amaré, *NSMB* (2015)). Values are *mean* ± *standard deviation* (*s.d.*) with *n* ≥ 3 independent titrations.

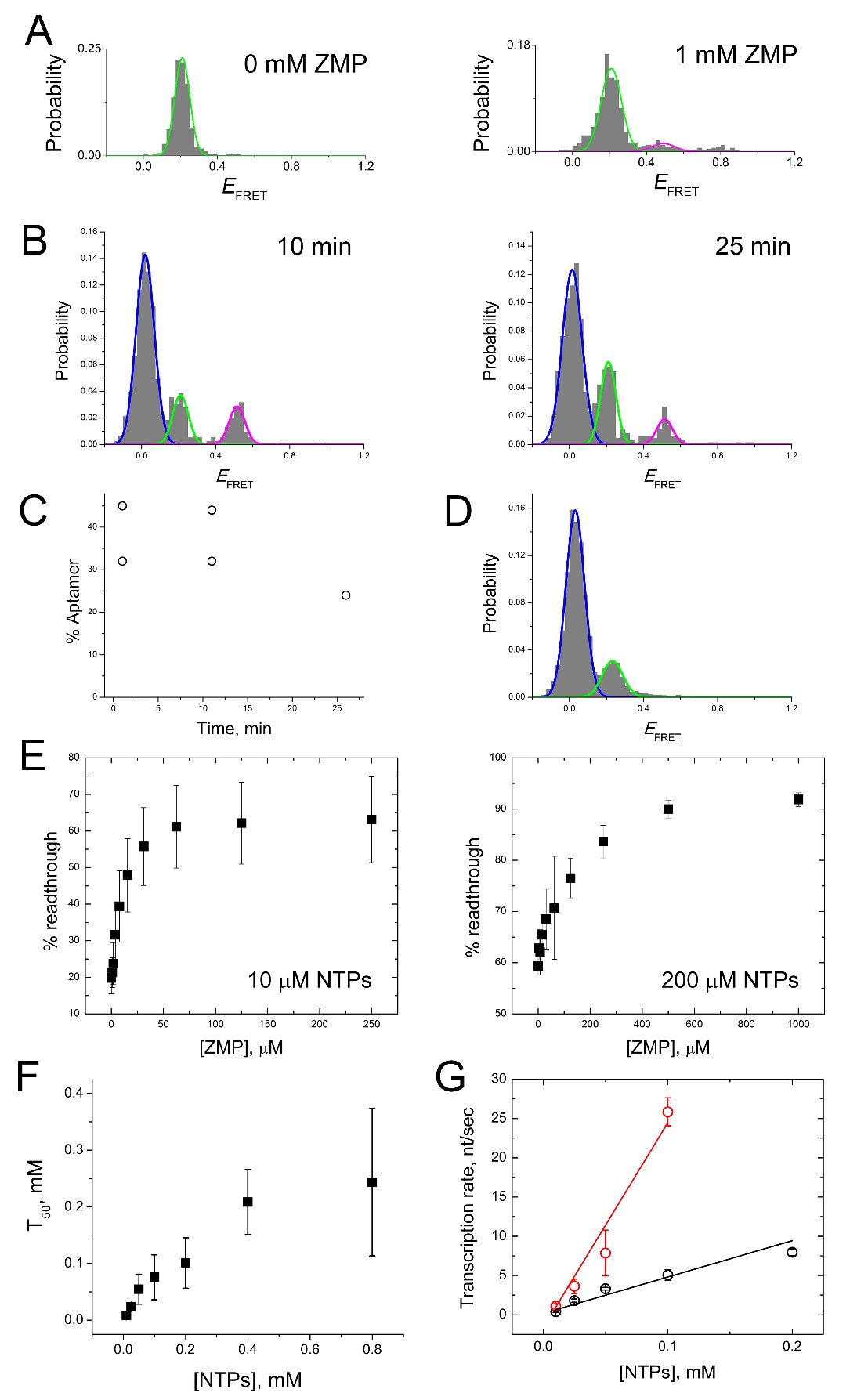

**Supplementary Figure S2 (Related to Figure 3)** Single-molecule VF assays and bulk transcription experiments of WT ZTP riboswitch. (**a**) *E*_FRET_ histograms of thermally refolded WT ZTP riboswitch at 0 mM and 1 mM ZMP. (**b**) The *E*_FRET_ histograms of WT that was vectorially folded at 1 mM ZMP and incubated for 10 min and 25 min before being imaged. (**c**) The remaining percentage of aptamer *vs*. the post-ATP addition time in VF assays. The percentage of aptamer is calculated as the ratio of the aptamer population (magenta) over the sum of the terminator (green) and aptamer populations. (**d**) The *E*_FRET_ histogram of WT that was vectorially folded for 30 s at 0 mM ZMP before 1 mM ZMP was added to stabilize the aptamer conformation. The 30 s was shortened from a longer 1-min folding time that was used in other VF experiments. (**e**) Bulk single-round transcription termination experiments at 0.01 mM NTPs (left) and 0.2 mM NTPs (right) are shown. Values are *mean* ± *s.d.* with *n* ≥ 3 independent experiments. The transcription midpoints (T50) are determined from fits to these titrations. (**f**) The T50 values from bulk single-round transcription termination experiments are shown at different NTP concentrations (*mean* ± *s.d.*, *n* ≥ 3 independent experiments). (**g**) The apparent rates of RNA synthesis for the terminated (red) and readthrough (black) transcription products are shown at different NTP concentrations (*mean* ± *s.d.*, *n* ≥ 3 independent experiments).

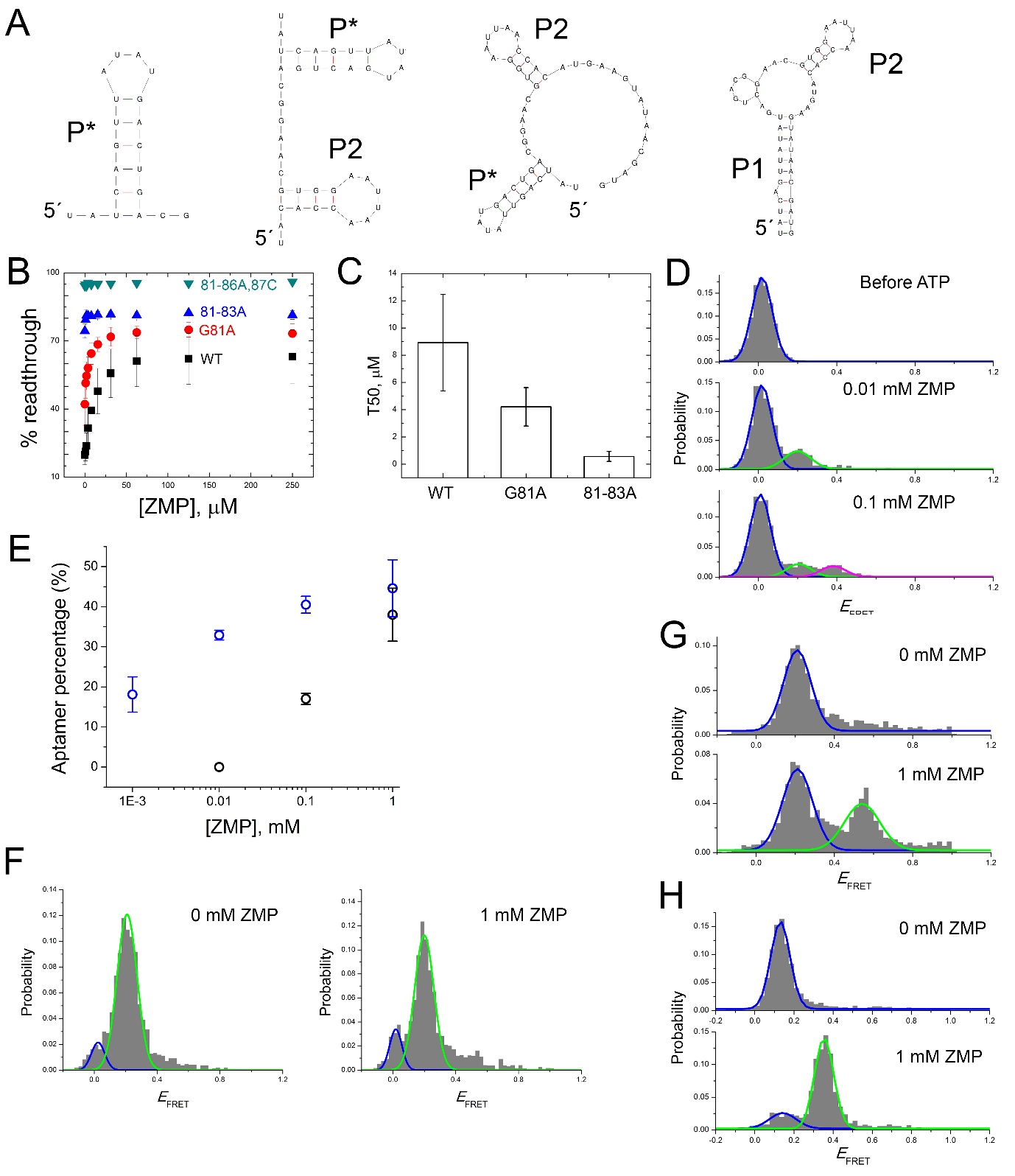

**Supplementary Figure S3 (Related to Figure 4)** Single molecule helicase unwinding assays and bulk single-round transcription termination experiments with ZTP riboswitch terminator variants and pause mimics. (**a**) Sequential *in silico* folding of the ZTP riboswitch. From left to right, the predicted secondary structure of nt 1-20, 1-40, 1-53, and 1-53 (alternative fold) of the WT ZTP riboswitch sequence. The helix formed by nt 3-8 and 13-18 is labeled P* and is not found in the aptamer fold (Fig. 1). The alternative fold for nt 1-53 contains helices P1 and P2, which are found in the aptamer fold. (**b**) ZTP riboswitch variants containing terminator mutations are titrated with ZMP in single round transcription experiments. Values are *mean* ± *s.d.* with *n* ≥ 3 independent experiments. (**c**) T50 values are shown for the terminator variants in **b**. The 81-86A,87C variant was not fit as no change was apparent over this concentration range. Values are *mean* ± *s.d.* with *n* ≥ 3 independent experiments. (**d**) *E*_FRET_ histograms of the 81-83A heteroduplex before and after ATP addition at different ZMP concentrations, as measured by VF with Rep-X. (**e**) The percentage of aptamer fold obtained by VF by Rep-X for WT and terminator mutant 81-83A in the presence of ZMP (*mean* ± *s.e.m.*, *n* = 3-4). (**f**) The *E*_FRET_ histograms of refolded 81–83A at 0 mM and 1 mM ZMP. To distinguish the terminator from the ZMP-unbound aptamer, 81–83A was imaged at 1 mM ZMP after being refolded at different ZMP concentrations. (**g**) *E*_FRET_ histograms of terminator mutant 92-94A, thermally refolded and imaged in the absence and presence of 1 mM ZMP. (**h**) *E*_FRET_ histograms of terminator mutant 81-83,92-94A, thermally refolded and imaged in the absence and presence of 1 mM ZMP.

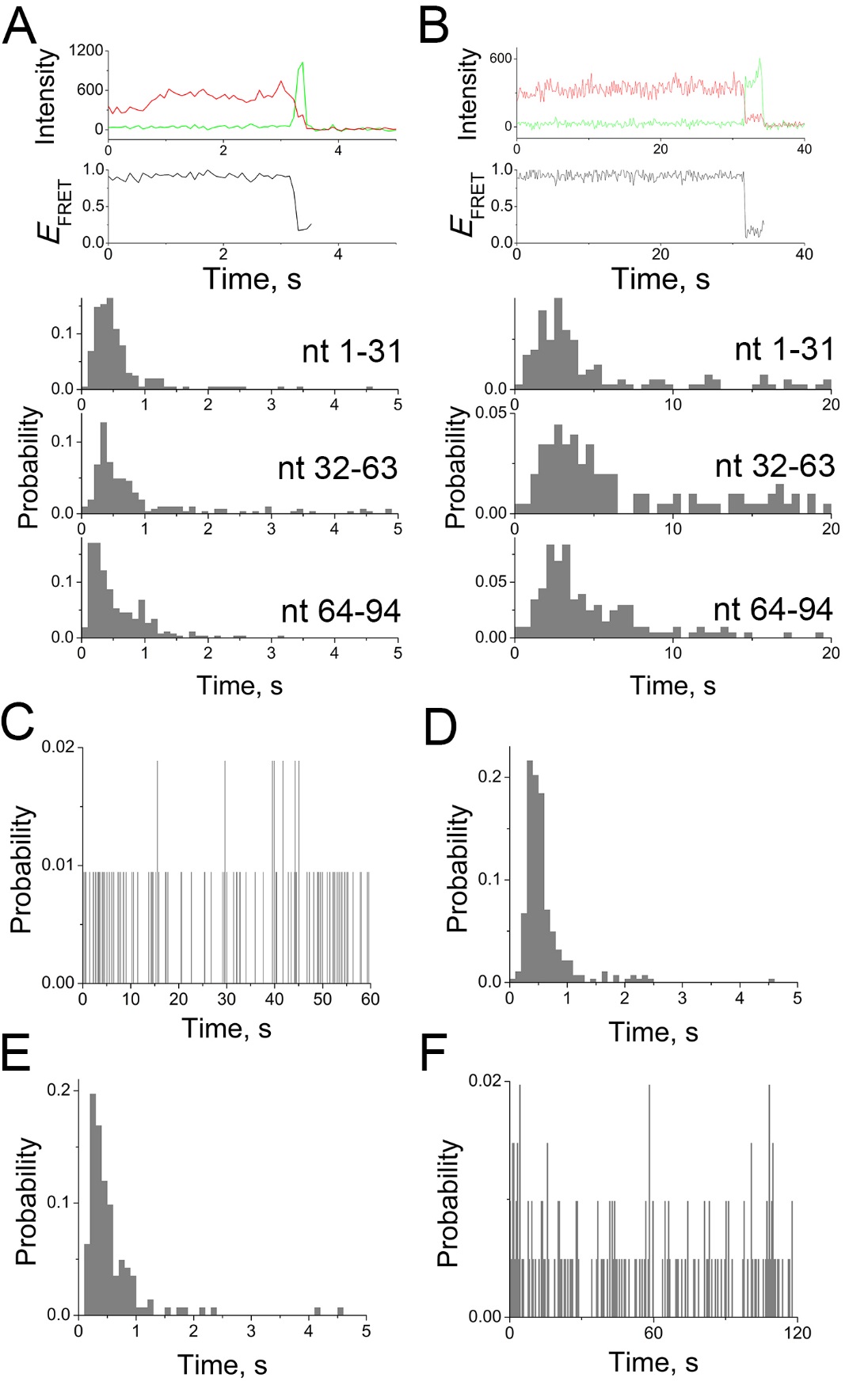

**Supplementary Figure S4 (Related to Figure 5)** Helicase unwinding rate via dropoff measurements (**a**) Individual single molecule trajectory (top) and histogram of all dropoff unwinding events (bottom) for Rep-X unwinding of indicated segments of the ZTP riboswitch. (**b**) Individual single molecule trajectory (top) and histogram of all dropoff unwinding events (bottom) for PcrA-X unwinding of indicated segments of the ZTP riboswitch. Intensity dropoff experiments to measure helicase unwinding rate. See Figure 4a (middle) for overall experimental design, in which unwinding causes a loss in FRET followed by a loss in Cy3 fluorescence as labeled oligos dissociate after unwinding. (**c**) Observation of photobleaching of hybrid duplex in the absence of ATP for Rep-X. (**d**) Histograms of time differences (Δt) for all molecules for a Cy5-labeled DNA oligo and Cy3-labeled RNA oligo. (**e**) Histograms of Δt for all molecules for a Cy5-labeled RNA oligo and Cy3-labeled DNA oligo. (**f**) Observation of photobleaching of hybrid duplex in the absence of ATP for PcrA-X.

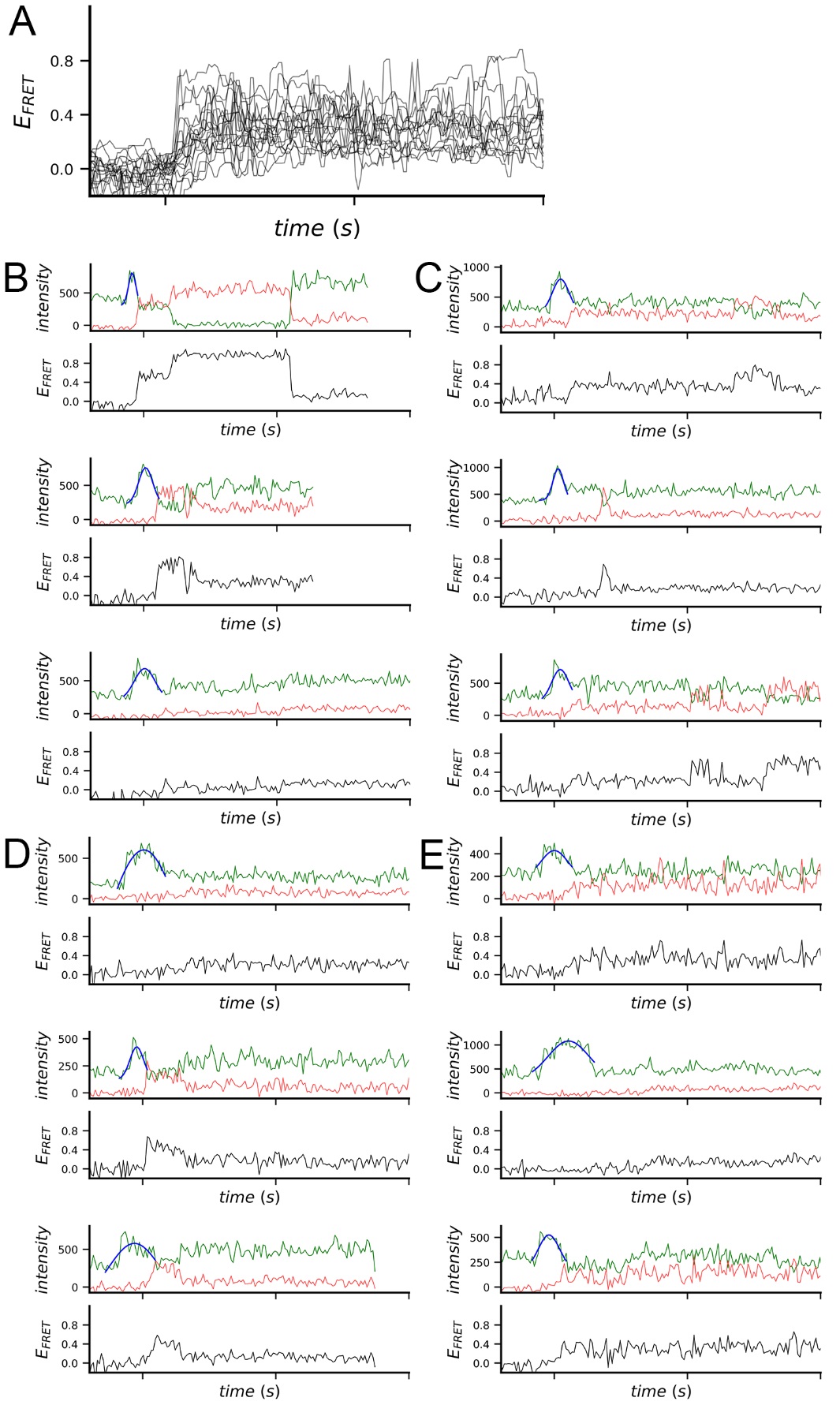

**Supplementary Figure S5 (Related to Figure 6)** Real-time unwinding measurements example trajectories. (**a**) Real-time trajectories of terminator mutant 81–83A obtained at 1 mM ZMP in VF assays using Rep-X. Shown are the overlay of all 81–83A (n = 19) trajectories (bottom) obtained at 1 mM ZMP. Trajectories are synchronized according to the PIFE peak center at position U32. The time duration between ticks is 5 s. (**b**) Example real-time trajectories of WT, and (**c**) terminator mutant 81–83A using VF assays with Rep-X in the absence ZMP. (**d**) Example real-time trajectories of WT and (**e**) 81-83A using VF with Rep-X in the presence of 1 mM ZMP.

**Supplementary Table 1** **(Related to Figure 1)**

RNA and DNA oligonucleotide sequences used for making the single-molecule fluorescence constructs.

| **Name** | **Sequences** |
| --- | --- |
| Ligated *F. ulcerans* ZTP riboswitch (WT) | UAUCAGUUAUAUGACUGACGGAACGUGGAAUUAACCACA UGAAGUAUAACGAUGACAAUGCCGACCGUCUGGGCGAAC  AAGCCUAGAUUGUCGGUUCAUUAGCGGUAUUCCGGAAUU GCCGUAAUCGCG |
| Ligated *F. ulcerans* ZTP aptamer (Δterm) | UAUCAGUUAUAUGACUGACGGAACGUGGAAUUAACCACA UGAAGUAUAACGAUGACAAUGCCGACCGUCUGGGCGUUC AUUAGCGGUAUUCCGGAAUUGCCGUAAUCGCG |
| Ligated *F. ulcerans* ZTP riboswitch (81-83A) | UAUCAGUUAUAUGACUGACGGAACGUGGAAUUAACCACA UGAAGUAUAACGAUGACAAUGCCGACCGUCUGGGCGAAC AAAAAUAGAUUGUCGGUUCAUUAGCGGUAUUCCGGAAUU GCCGUAAUCGCG |
| Ligated *F. ulcerans* ZTP riboswitch (92-94A) | UAUCAGUUAUAUGACUGACGGAACGUGGAAUUAACCACA UGAAGUAUAACGAUGACAAUGCCGACCGUCUGGGCGAAC  AAGCCUAGAUUGUAAAUUCAUUAGCGGUAUUCCGGAAUU GCCGUAAUCGCG |
| Ligated *F. ulcerans* ZTP riboswitch (81-83, 92-94A) | UAUCAGUUAUAUGACUGACGGAACGUGGAAUUAACCACA UGAAGUAUAACGAUGACAAUGCCGACCGUCUGGGCGAAC  AAAAAUAGAUUGUAAAUUCAUUAGCGGUAUUCCGGAAUU GCCGUAAUCGCG |
| nt 1-33, Cy3 at U32 | UAUCAGUUAUAUGACUGACGGAACGUGGAAU-Cy3-UA |
| nt 1-33, Cy3 at U12 | UAUCAGUUA-Cy3-UAUGACUGACGGAACGUGGAAUUA |
| nt 34-71 | ACCACAUGAAGUAUAACGAUGACAAUGCCGACCGUCUG |
| nt 72-94, Cy3 at U84 | GGCGAACAAGCC-Cy3-UAGAUUGUCGGTTCATTAGCGGTA TTCCGGAATTGCCGTAAT CGCG |
| 18-speedmer | GCCUCGCUGCCGUCGCCA-Cy3 |
| biotin-DNA-Cy5 | biotin-CGCGATTACGGCAATTCCGGAATACCGCTAATG-Cy5 |
| dT_20_ cDNA to WT | CCGACAATCTAGGCTTGTTCGCCCAGACGGTCGGCATTGT CATCGTTATACTTCATGTGGTTAATTCCACGTTCCGTCAGT CATATAACTGATATTTTTTTTTTTTTTTTTTTT |
| dT_20_ cDNA to Δterm | CGCCCAGACGGTCGGCATTGTCATCGTTATACTTCATGTGG TTAATTCCACGTTCCGTCAGTCATATAACTGATA TTTTTTTTTTTTTTTTTTTT |
| dT_20_ cDNA to 81-83A | CCGACAATCTATTTTTGTTCGCCCAGACGGTCGGCATTGTC ATCGTTATACTTCATGTGGTTAATTCCACGTTCCGTCAGTC ATATAACTGATATTTTTTTTTTTTTTTTTTTT |
| dT_30_ short cDNA to nt 56-94 | CCGACAATCTAGGCTTGTTCGCCCAGACGGTCGGCATTG TTTTTTTTTTTTTTTTTTTTTTTTTTTTTT |
| cDNA-speedmer-dT_20_ | CCGACAATCTAGGCTTGTTCGCCCAGACGGTCGGCAT TGTCATCGTTATACTTCATGTGGTTAATTCCACGTTCC GTCAGTCATATAACTGATATTTGGCGACGGCAGCGAG GCTTTTTTTTTTTTTTTTTTTT |
| *F. ulcerans* ZTP aptamer plasmid^1^ | TAATACGACTCACTATAGGGAGATATAACTGATACTGAT GAGTCCGTGAGGACGAAACGGTACCCGGTACCGTCTAT CAGTTATATGACTGACGGAACGTGGAATTAACCACATGA AGTATAACGATGACAATGCCGACCGTCTGGGCG |
| *F. ulcerans* ZTP transcription termination plasmid^1^ | TTGACTATTTTACCTCTGGCGGTGATAATGGTTGCAATG TAGTAAGGAGGTTGTATGGAAGATTATCAGTTATATGAC TGACGGAACGTGGAATTAACCACATGAAGTATAACGAT GACAATGCCGACCGTCTGGGCGAACAAGCCTAGATTGTC GGTTTTTTTTATACATTTTTTTAGGAGGAGATAATAAGGG AATATCAAATATAATTGTTGA |
| *F. ulcerans* ZTP transcription termination plasmid G81A^1^ | TTGACTATTTTACCTCTGGCGGTGATAATGGTTGCAATG TAGTAAGGAGGTTGTATGGAAGATTATCAGTTATATGAC TGACGGAACGTGGAATTAACCACATGAAGTATAACGAT GACAATGCCGACCGTCTGGGCGAACAAACCTAGATTGTC GGTTTTTTTTATACATTTTTTTAGGAGGAGATAATAAGGG AATATCAAATATAATTGTTGA |
| *F. ulcerans* ZTP transcription termination plasmid 81-83A^1^ | TTGACTATTTTACCTCTGGCGGTGATAATGGTTGCAATG TAGTAAGGAGGTTGTATGGAAGATTATCAGTTATATGAC TGACGGAACGTGGAATTAACCACATGAAGTATAACGAT GACAATGCCGACCGTCTGGGCGAACAAAAATAGATTGTC GGTTTTTTTTATACATTTTTTTAGGAGGAGATAATAAGGG AATATCAAATATAATTGTTGA |
| *F. ulcerans* ZTP transcription termination plasmid 81-86A,87C^1^ | TTGACTATTTTACCTCTGGCGGTGATAATGGTTGCAATG TAGTAAGGAGGTTGTATGGAAGATTATCAGTTATATGAC TGACGGAACGTGGAATTAACCACATGAAGTATAACGAT GACAATGCCGACCGTCTGGGCGAACAAAAAAAAACTGTC GGTTTTTTTTATACATTTTTTTAGGAGGAGATAATAAGGG AATATCAAATATAATTGTTGA |
| Primers to delete nt 1-33 from the plasmid | ACGGTACCCGGTACCGTCACCACATGAAGTATAACG, CGTTATACTTCATGTGGTGACGGTACCGGGTACCGT |
| Primers to change hammerhead in Δ1-33 plasmid | CGACTCACTATAGGGAGACTTCATGTGGTCTGATGAGT CCGTGAGGAC, GTCCTCACGGACTCATCAGACCACATGAAGTCTCCCTATA GTGAGTCG |
| PCR T7 promoter forward primer | TAATACGACTCACTATAG |
| PCR reverse primer to make Δ1-33 with tether RNA for WT | mCmGCGATTACGGCAATTCCGGAATACCGCTAATGAACC GACAATCTAGGCTTGTTCGCCCAGACGGTCGGCAT^2^ |
| PCR reverse primer to make Δ1–33 with tether RNA for Δterm | mGmCGATTACGGCAATTCCGGAATACCGCT  AATGAACGCCCAGACGGTCGGCAT^2^ |
| PCR reverse primer to make Δ1-33 with tether RNA for 81-83A | mCmGCGATTACGGCAATTCCGGAATACCGCTA ATGAACCGACAATCTATTTTTGTTCGCCCAGAC GGTCGGCAT^2^ |
| PCR reverse primer to make Δ1-33 with tether RNA for 92-94A | mCmGCGATTACGGCAATTCCGGAATACCGCTA ATGAATTTACAATCTAGGCTTGTTCGCCCAGAC GGTCGGCAT^2^ |
| PCR reverse primer to make Δ1-33 with tether RNA for 81-83, 92-94A | mCmGCGATTACGGCAATTCCGGAATACCGCTA ATGAATTTACAATCTATTTTTGTTCGCCCAGAC GGTCGGCAT^2^ |

Notes:

^1^For plasmids, only the promoter and transcribed regions are shown.

^2^Nucleotides with 2´-O-methyl modifications are preceded by “m”.
